# Regulation of odd chain fatty acid metabolism in the development of metabolic diseases in mice fed a low protein diet

**DOI:** 10.1101/2022.05.27.493394

**Authors:** Isaac Ampong

**Affiliations:** Center for Precision Medicine, Wake Forest University Baptist Medical Center, Medical Center Boulevard, Winston-Salem, North Carolina, United States

**Keywords:** low protein diet, OCFA, gene expression, metabolism, metabolic diseases

## Abstract

Nonalcoholic fatty liver disease (NAFLD) and Metabolic syndrome (MS) have become a global health concern as incidence of these metabolic disorders is growing rapidly in developing countries particularly in the Middle East, South America and Africa. Studies have shown that protein restriction is associated with increased risk of metabolic diseases, possibly through effects on fatty acid (FA) metabolism. In the present study, we investigated whether a low protein diet modulates FA metabolism and whether methyl donor supplementation can ameliorate these effects and improve metabolic health. Male C57BL/6 mice were fed either a low protein diet (LPD, 90 g/kg protein, n=8), a LPD supplemented with methyl donors (MD-LPD; choline chloride, betaine, methionine, folic acid, vitamin B12, n=8) or normal protein diet (NPD, 180 g/kg protein, n=8) for 7 weeks prior to analysis of serum fatty acid profiles by GC FID and MS and liver fatty acid synthesis and uptake gene expression by RT-qPCR. We observed significant depletion of serum C15:0 and C17:0 in LPD-fed males compared to NPD. Serum long chain saturated FAs C18:0 and C24:0 were increased in LPD male mice compared to NPD. Gene expression analysis revealed an upregulation of hepatic cluster of differentiation 36 (*CD36)* expression in LPD mice compared to NPD suggesting increased fat uptake in the liver. However, when LPD diet was supplemented with methyl donors, we observed either no change in serum C15: 0 and an increased serum C17:0 compared to LPD with no methyl donor supplementation. Again, methyl donor supplementation upregulated fatty acid desaturase 1 (*FADS1)*, thioredoxin-1 *(TRX1)* and catalase *(CAT)* expression in the liver of MD-LPD fed mice compared to LPD mice. Altogether, our study revealed that odd chain fatty acids (OCFA)s are key early markers observed in a suboptimal diet-induced metabolic changes and may be potential targets to improve metabolic health outcomes.

## 1. Introduction

Metabolic syndrome (MS) is closely associated with an increased risk of chronic diseases such as type 2 diabetes and cardiovascular disease which are leading cause of mortality globally **Lin et al (2020)**. Furthermore, several cross-sectional studies have linked NAFLD to diabetes and MS suggesting NAFLD precedes the development of these conditions **Lonardo et al (2015)**. Lipids play crucial roles in the pathogenesis of various metabolic conditions such as NAFLD, insulin resistance and MS **Denisenko et al (2020)**. Accumulated evidence suggests that different FAs play specific roles depending on their carbon length, degree of saturation, chain conformation and whether they are an even or odd chain FA **Zhou et al (2020)**. For instance, excessive consumption of saturated fats contributes to the pathogenesis of obesity and related diseases. It has also been shown that high plasma levels of free fatty acids (FFAs), particularly saturated fatty acids (SFAs), are associated with insulin resistance in obese patients with type 2 diabetes mellitus **Denisenko et al (2020)**. However, in recent times, OCFA have been suggested as important targets to improve metabolic diseases in part due to their inverse association with several cardiometabolic events. In addition to the gut microbiota and diet suggested as potential source of OCFA, several metabolic pathways including phytanoyl-CoA α-hydroxylase *(PHYH)*, 2-hydroxyphytanoyl-CoA lyase *(HACL1)*; pristanal dehydrogenase, branched-chain alpha-keto acid dehydrogenase *(BCKDHA)* are known to be involved in their synthesis **Ampong et al (2021)**.

Proteins are an important metabolic fuel source. However, since excess proteins cannot be stored directly in the body, they must be converted into glucose or triglycerides. The liver is a major regulatory organ for protein and amino acid metabolism. In endogenous protein metabolism, the liver is essential for the formation of plasma proteins (eg albumin, clotting factors), amino acid interconversion (synthesis of all non-essential amino acids), deamination of amino acids to produce energy and urea synthesis (for ammonia excretion), **Charlton (1996)**. NAFLD, which is a precursor for the development of T2DM and MS, is highly prevalent in the developing world particularly in the Middle East, South America and Africa. Studies in rodents and humans have shown that a low-protein diet (generally defined as < 9 % of total energy) causes several metabolic disturbances including increased oxidative stress, hepatic steatosis, glucose intolerance as well as elevated adiposity **Kuwahata et al (2011), Kang et al (2011), Jahan-Mihan et al (2015)**.

Methionine is an amino acid required for protein synthesis and is a constituent of the tripeptide antioxidant, glutathione. It has been reported that methyl donor deficiency induces liver steatosis and predisposes to MS, **Pooya et al (2012)**. Methyl donor supplementation reverted the high fat sucrose (HFS)-diet-induced hepatic TG accumulation in rats after 8 weeks of feeding **Cordero et al (2013)**. Although few studies have investigated the development of MS in animals fed a low protein diets **Kuwahata et al (2011), Kang et al (2011), Jahan-Mihan et al (2015)**, however, these studies have largely focused on the role of free fatty acid (FFA) in NAFLD rather than on OCFA. Therefore, we proposed that protein restricted dietary consumption may decrease systemic OCFA metabolism and that methyl donors supplement in a protein restricted diet may improve OCFA metabolism and decrease cardiometabolic risks.

## 2. Materials and Methods

### 2.1 Experimental Diets and Animals

Eight-week-old male C57BL/6 mice were purchased from Harlan Ltd, Belton, Leicestershire, UK and were maintained in a temperature (20±2 ^0^C) with a 12h day/light cycle. All experimental and study procedures were conducted under the UK Home Office Animal (Scientific Procedures) Act 1986 Amendment Regulations 2012, which transposed Directive 2010/63/EU into UK law, and were approved by the Animal Welfare and Ethical Review Board at Aston University.

After one week of acclimatization, animals were fed either control NPD; 18% casein; *n* = 8), LPD; 9% casein; *n* = 8 or MD-LPD; 5 g/kg diet choline chloride, 15 g/kg diet betaine, 7.5 g/kg diet methionine, 15 mg/kg diet folic acid, 1.5 mg/kg diet vitamin B12; *n* = 8). Diets were manufactured commercially (Special Dietary Services Ltd; UK) and their composition is shown in supplementary Table S1. Following the 7-weeks of feeding, all animals were fasted for 8 hours and sacrificed via cervical dislocation. Blood samples were taken via heart puncture, centrifuged at 10,600 ***g*** (4°C, 10 min) and the serum aliquoted and stored at −80°C. Liver samples were dissected and weighted prior to being snap frozen and stored at −80°C until used.

### 2.2 Serum and Liver fatty acid (FA) analysis

Serum FA extraction was determined using a methodology previously described by Pararasa et al (2016). Following serum thawing, a 2.63 μg/mL internal standard (Undecanoic acid; C11:0) (Sigma) was added to 50 μL of mouse serum prior to being diluted to a final volume of 450 μL in PBS (Thermo-Fisher). Fatty acids were extracted using a chloroform-methanol mix (2:1; Thermo-Fisher) in 0.01% butylated hydroxytoluene (Sigma) prior to centrifugation at 200 ***g*** for 10 min at 4°C. The organic (chloroform) phase was removed and dried under nitrogen gas. The isolated fatty acids were methylated (200 μL toluene, 0.3 mL (6.3%) HCL)) in 1.5 mL methanol (all from Thermo-Fisher) at 100°C for 1 h in PTFE-sealed glass vials. The fatty acid methyl esters (FAMEs) were subsequently extracted with 1 mL of hexane (Thermo- Fisher) and 1 mL of water, evaporated under nitrogen and re-suspended in 20 μL of hexane prior to analyses by gas chromatography (7820A (G4350A) GC system; Agilent technologies) equipped with a flame-ionization detector and using an Omegawax 250 capillary column (30 m × 0.25 mm ID × 0.25 µm film; Sigma). For liver sample preparation, 50mg of frozen tissue was thawed on ice and homogenized in 80µL normal saline and 2.63 μg/mL undecanoic acid (C11:0) was spiked in as an internal standard. Methylation of liver FA was conducted at 35 °C for 10 min and following same protocol as described above except that the re-suspended samples were analysed by GCMS performed at the Stable Isotope Laboratory, Leggett Building, Faculty of Health and Medical Sciences, University of Surrey, Guildford. Identification and quantification of FA peaks were determined against peak areas of a Supelco 37 FAME mix standard. In the GC-MS analyses, in addition to the internal standard (IS), a calibration curve was constructed for heptadecanoic acid (C17:0) against C11:0. A calibration plot of C17:0 compound was run by applying the ratio of the peak area of the FAME in the standards to the peak area of the IS against the ratio of the concentration of the FAME to the concentration of the IS. The concentration of FAME in the heptadecanoic acid (C17:0) solution was then determined using the area ratio and the calibration plot. The composition of the FAs (μM) in the samples was then recalculated and used to determine percentage fatty acid composition.

### 2.3 Gene expression analysis by qPCR

Total RNA of the liver was isolated using RNeasy Lipid Tissue Mini Kit (QIAGEN, UK) and the concentration and quality were determined by A260/280 ratios by Nanodrop Lite Spectrophotometer (Thermo Scientific, UK) prior to cDNA synthesis using the NanoScript (PrimerDesign, UK) kit, all according to the manufacturer’s instructions. Real-Time PCR (RT-qPCR) analysis was carried out using Stratagene (Thermo Scientific, UK) or Quant studio 7 Real-Time PCR system (Thermo Scientific, UK). RT-qPCR reactions for tissue samples were prepared using Precision SYBRgreen Mastermix (Primerdesign, UK), 175 nM forward and reverse primers (Eurofins) and water. A post-amplification melting curve confirmed the presence of specific products for each primer set. Ct values were converted to relative expression values using the delta-delta Ct method with gene-of-interest expression normalised to *Pgk1* and *Tbp* for hepatic tissue using geNorm software as described previously **(Lucas *et al*. 2011)** and NormFinder. The primer sequences for genes investigated, *CD36, FADS1, CAT, BCKDHA*, 2-hydroxyacyl-CoA lyase 1(*HACL1*), carnitine palmitoyltransferase 1b, muscle (*CPT1*), carnitine palmitoyltransferase 2(*CPT2*), tumor necrosis factor (*TNF- α*) and glutathione reductase *(GSR)* are provided in supplementary Table 1.

### 2.4 Statistical analyses

All statistical analyses were performed using the Prism vs 8 (GraphPad, UK) and data normality assessed with Shaprio–Wilk and Kolmogorov–Smirnov tests. Data were expressed as percentages and mean (standard deviation) as appropriate for normally distributed data, or median (range) for non-normally distributed data. Statistically significance by t-test, or Kruskal–Wallis test with Dunns multiple comparison test was taken at *P* < 0.05.

## 3.0 Results

### 3.1 Body weight

During the experimental period, eight-week-old C57BL/6 males were housed singly and fed three different diets either control normal protein diet (NPD; n = 8) isocaloric low protein diet (LPD; n = 8) or MD-LPD; n=8) *ad libitum* for 7 weeks (Table 2.1). The NPD (controls) or LPD were exhibited a significant decrease in the body weight compared to MD-LPD fed mice (*p* < 0.05; n=8 mice) from week 5 to the day they were sacrificed (week 7). (Figure 1a). There was no difference between LPD and MD-LPD groups.

**Figure 1:**
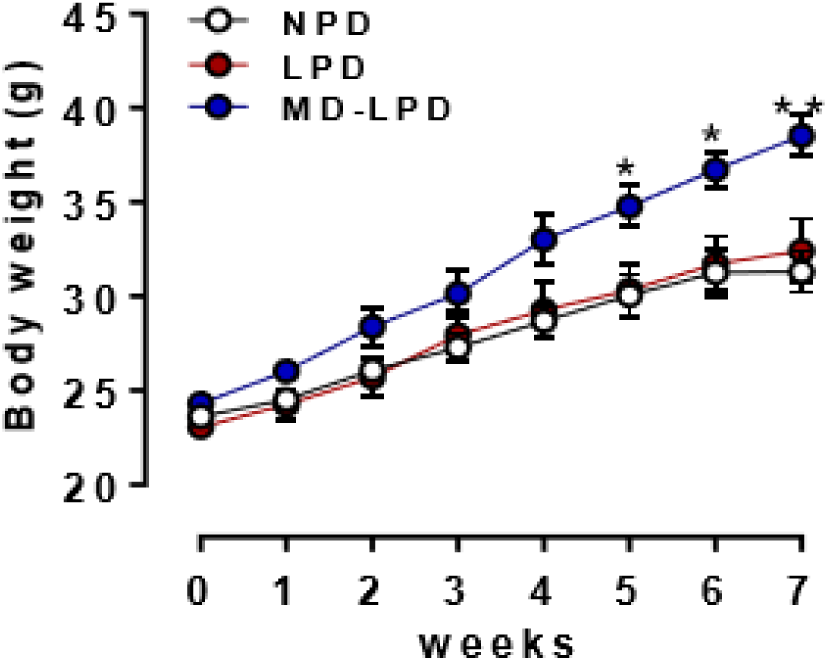
Body weight changes with time. Data represent mean ± SEM. Weekly body weight of male mice fed either a normal-protein diet (NPD; 18% casein; n=8), low-protein diet (LPD; 9% casein; n=8) or LPD supplemented with methyl donors (MD-LPD; 9% casein with 5 g/kg choline chloride, 15 g/kg betaine, 7.5 g/kg methionine, 15 mg/kg folic acid, 1.5 mg/kg vitamin B12; n=8). *p<0.05; **p<0.001

### 3.2 FA analysis of LPD in comparison between the NPD and MD-LPD groups

To explore the effect of dietary low protein intake on OCFA metabolism, we analysed serum and liver samples obtained from male mice fed LPD for 7 weeks. While no differences were observed in the liver total OCFA proportion between dietary groups, serum total OCFAs were significantly lower in LPD fed group [4.14 (3.04-6.03)] compared to NPD group [7.08 (5.21-10.56), p<0.05], and serum total OCFAs were significantly higher after MD-LPD [9.94 (8.24-11.54)] relative to LPD fed group [4.14 (3.04-6.03), p<0.001]. The proportion of serum either C17:0 or C15:0 was significantly lower in LPD animals compared to NPD (p<0.05). However, the proportion of serum C17:0 in MD-LPD mice was significantly higher when compared to those fed with LPD (figure 2, p<0.001).

**Figure 2:**
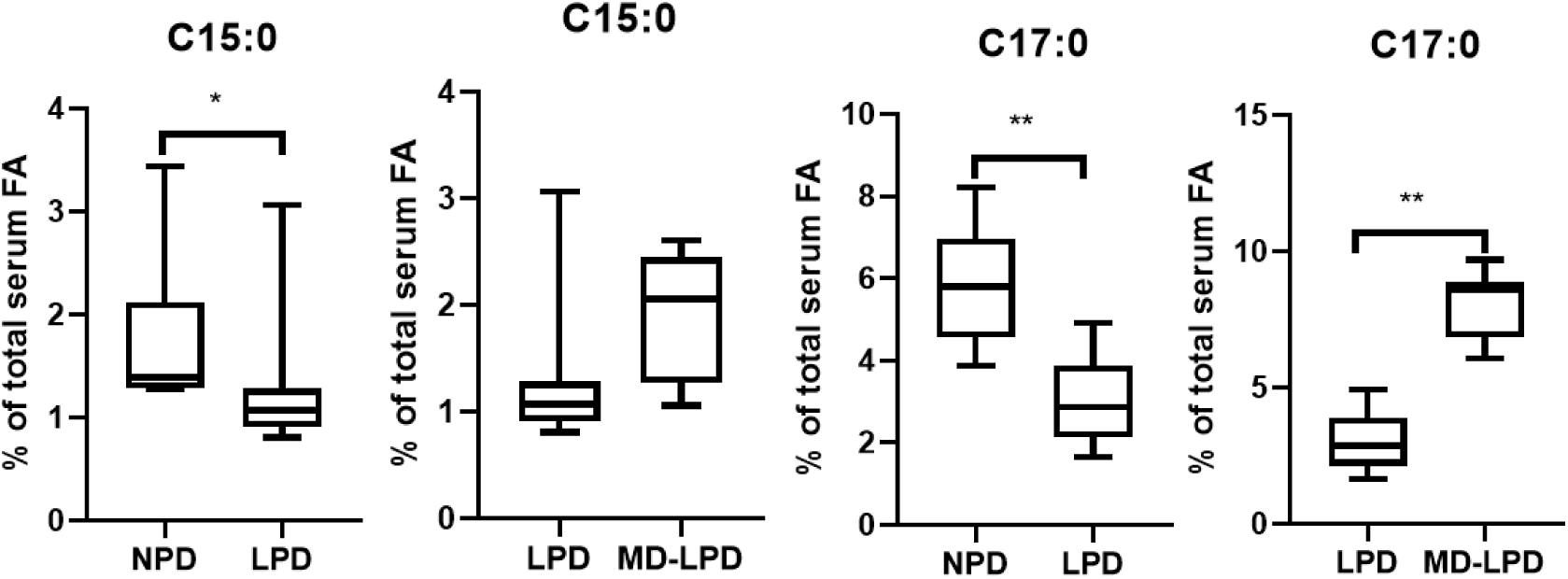
Effect of low protein intake on odd-chain saturated FA in the serum (Aston LPD study) Specific OCFAs was measured in the serum and liver of mice. Values are given as means ± SEM for n=8. *p<0.05 LPD vs NPD; **p<0.001 MD-LPD vs LPD

The percentage of total even-chain SFAs in serum were not significantly different in LPD-fed mice [43.89(43.57-65.21)] when compared to NPD-fed group [53.59 (49.29-57.50)], however, the MD- LPD-fed group [56.23(52.76-.21)] showed significantly higher even-chain SFAs compared to the LPD group [43.89(43.57-65.21), p<0.001]. In contrast, the proportion of liver total even-chain SFAs was significantly reduced in LPD [35.77(27.89-43.07)] relative to NPD [41.82(33.02- 56.42), p<0.05]. The addition of MD to the LPD [34.81(31.46-41.77)] had no effect on liver even-chain SFA proportion when compared to the LPD fed group [35.77(27.89-43.07)]. LPD group showed a lower concentrations of serum C10:0, C14:0 and C16:0 (p<0.05) and an increased concentrations of serum C18:0 (p<0.05) and C24:0 (p<0.001), compared to NPD (figure 3) whereas, MD-LPD group showed an increased proportion of serum C10:0 (p<0.001), C12:0 (p<0.05) and C14:0 (p<0.001) but decrease C24:0 (p<0.0001) compared to LPD (figure 3). In contrast, hepatic C18:0 (p<0.001) concentration was significantly lower in the LPD fed mice compared to NPD group (figure 3). There was no significant difference in liver C14:0 and C16:0 concentration between NPD and mice fed the LPD. Similar to serum MD-LPD levels of C14:0 (p<0.001), there was an increase concentration of hepatic C14:0 (p<0.05) when compared to LPD fed mice.

**Figure 3:**
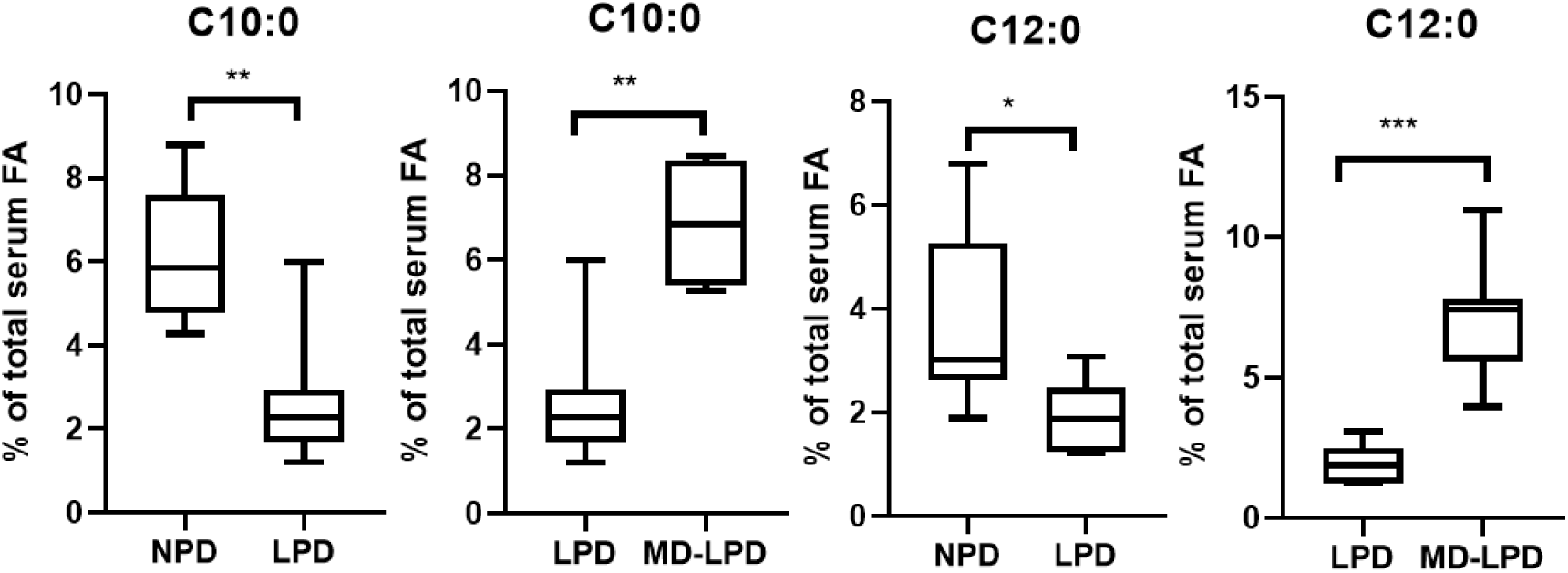
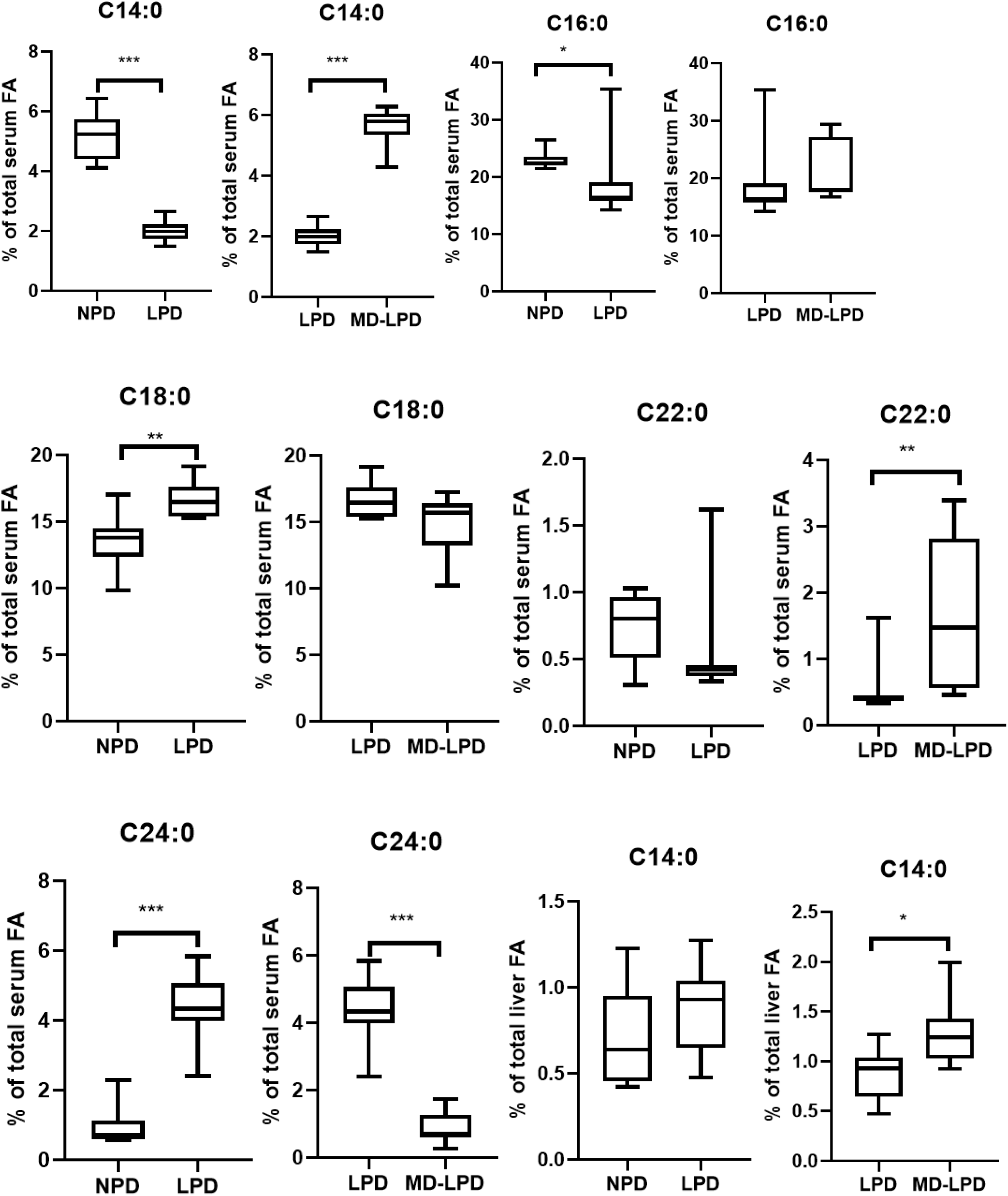
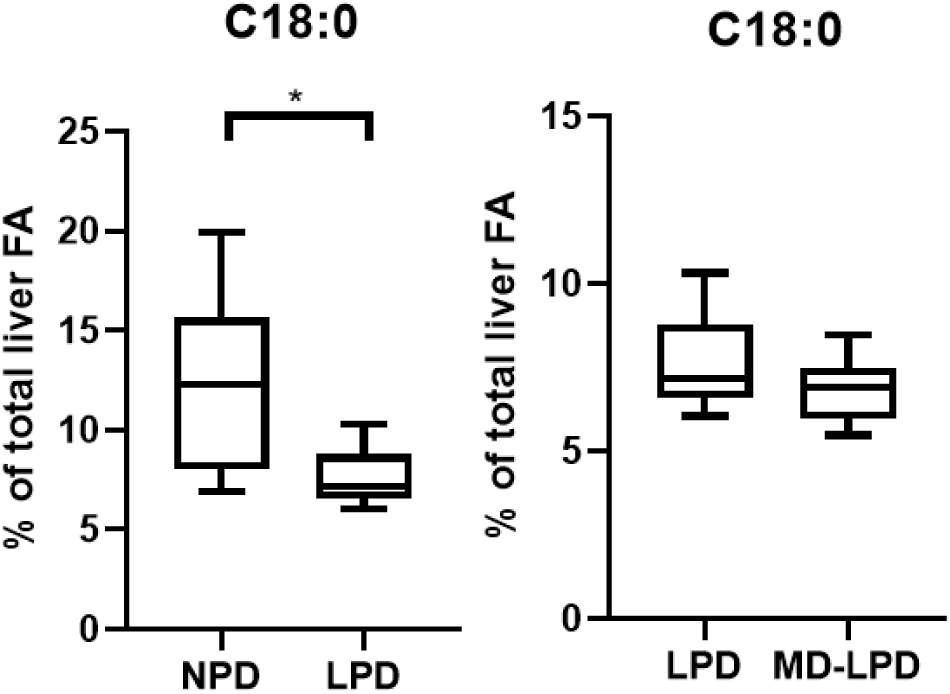
Effect of low protein intake on even-chain saturated FA in the serum and liver. Specific ECFAs were measured in the serum and liver of mice. Values are given as means ± SEM for n=8. *p<0.05, **p<0.001, ***p<0.0001 LPD vs NPD; *p<0.05, **p<0.001, ***p<0.0001 MD- LPD vs LPD

Total MUFA in either serum or liver was not significantly different between dietary groups. Serum content of palmitoleic acid (C16:1 FA) was significantly reduced in LPD fed mice than NPD group (p<0.05), however, they were reversed when LPD was compared with MD-LPD in the serum (figure 4, p<0.05). On the contrary, whilst serum proportion of oleic acid (C18:1 FA) was significantly lower in MD-LPD relative to LPD (p<0.05), there was no change when LPD was compared with NPD (Figure 4). Interestingly, individual MUFA such as palmitoleic acid (C16:1) and oleic acid (C18:1) in the liver of animals fed either NPD, LPD or MD-LPD did not differ between groups (figure 4).

**Figure 4:**
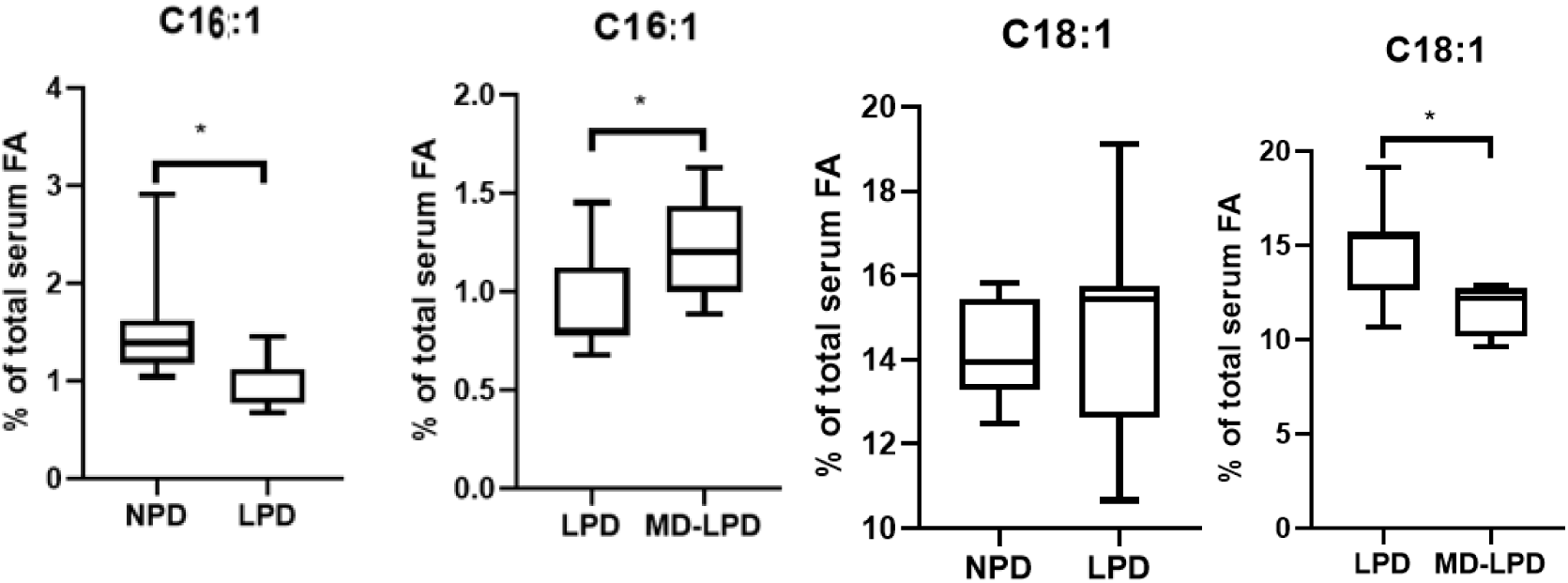
Effect of low protein intake on monounsaturated FA content in the serum and liver. Specific MUFAs were measured in the serum and liver of mice. Values are given as means ± SEM for n=8. *p<0.05 LPD vs NPD; *p<0.05 MD-LPD vs LPD)

Total serum n-6 PUFA content was significantly lower in MD-LPD [20.00(7.87-25.30)] compared to LPD [35.16(17.96-36.42), p<0.001]. In the liver, a lower but insignificant levels of total n-6 PUFA in MD-LPD fed mice [22.07(18.55-26.98)] compared to LPD without methyl donors supplementation 28.04(24.08-40.86). However, no observable change were recorded in both serum and liver total n-6 PUFA between LPD and NPD. The serum proportion of C20:4n6 FA was significantly higher in LPD compared to NPD (p<0.001), however, this was lowered when MD-LPD was compared with LPD. (Figure 5). Conversely, serum C22:2 levels were significantly lower in LPD group compared to NPD (p<0.05), whilst no significant difference was obtained between MD-LPD and LPD (Figure 5.5). In terms of serum content of C18:2n6, analyses of intergroup differences showed that there were significantly lower levels in MD-LPD compared to LPD (p<0.05), although no significant difference was seen when LPD fed mice were compared with NPD. (Figure 5). Despite changes in specific PUFAs in serum, liver levels of C18:2n6, C18:3n3 and C20:4n6 were not significant when LPD is compared to NPD fed mice. MD-LPD fed mice showed increased level of C18:2n6 compared to LPD mice in the liver.

**Figure 5:**
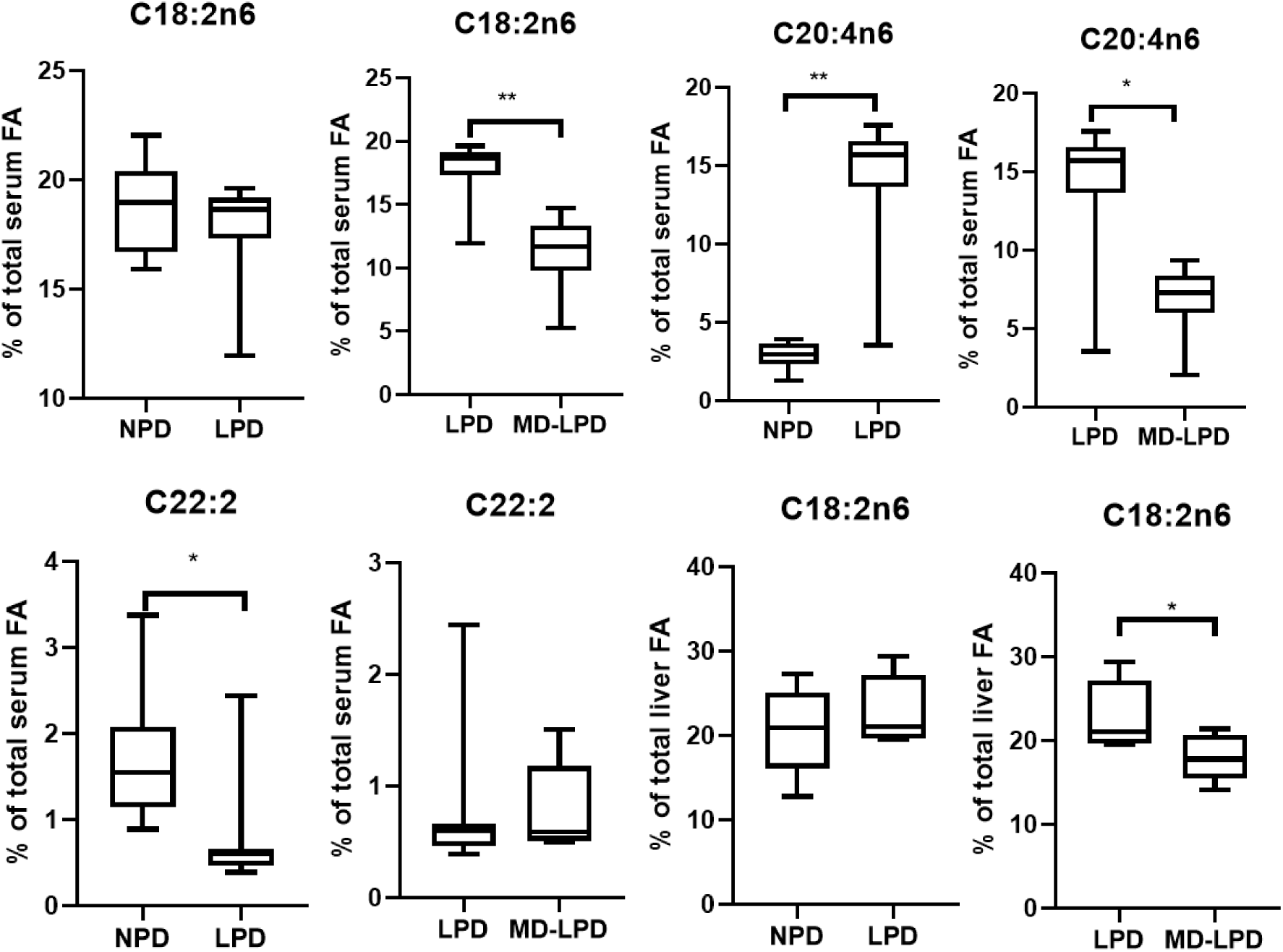
Effect of low protein intake on polyunsaturated FA content in the serum and liver. Specific PUFAs were measured in the serum and liver of mice. Values are given as means ± SEM for n=8. *p<0.05, **p<0.001 LPD vs NPD; *p<0.05 MD-LPD vs LPD

### 3.3 Gene expression profile among three experimental groups

Following a low protein dietary intake, gene expression analyses on key genes involved in hepatic fatty acid transport and metabolism, oxidative stress and inflammation were investigated using RT-PCR. There was significant upregulation of *CD 36* expression in the liver of LPD compared to NPD group (p<0.001), however, there were no significant changes in the mRNA expressions of *FADS1, HACL-1, BCKDHA, CPT1, CPT2, TNF, TRX1, GSR* and *CAT*. Comparing MD-LPD to LPD group, we observed that, *FADS1, TRX1* and *CAT* expressions were significantly higher in MD-LPD (p<0.001) although *CD 36, HACL-1* and *BCKDHA* expressions were not altered in MD- LPD (Figure 6).

**Figure 6:**
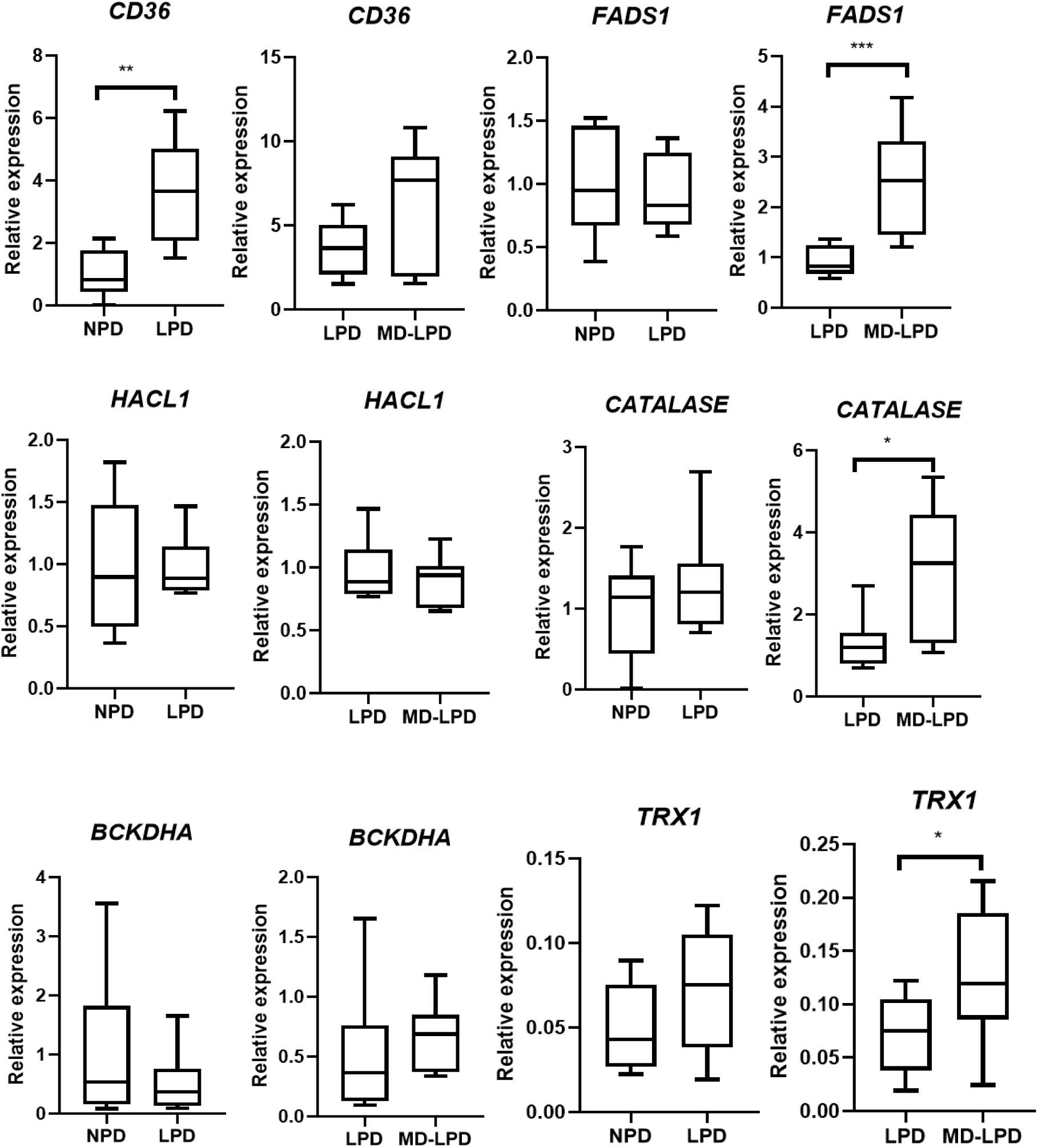
Effect of diet on mouse liver *CD 36, FADS1, CAT, TRX1, HACL-1* and *BCKDHA* mRNA expression. Mean relative transcript expression of genes related specific fatty acid changes in mouse. Values are given as means ± S.E.M for n=8; *p < 0.05, ** p < 0.001; ***p < 0.0001, NPD vs LPD or LPD vs MD-LPD.

## 4.0 Discussion

Dysregulation of fatty acid metabolism in the liver and serum appear to be the main factors underlying the pathogenesis of NAFLD and subsequent metabolic diseases development **Cortez-Pinto et al (2006); Yoo et al. 2017; Perdomo et al (2019)**. Although, a number of studies have linked an LPD with NAFLD and MS as evidence in in-vivo models studies where animals have been fed LPD in duration ranging between 5-8 weeks. There is limited evidence of this diet on specific fatty acid metabolism, particularly OCFA, which epidemiological evidence have inversely associated with type 2 diabetes and other cardiometabolic events Prada et al (2021). OCFAs, such as C15:0 and C17:0 have become important biomarkers in recent studies because of their consistent negative correlation with metabolic diseases. In several epidemiological studies, C15:0 and C17:0 have been shown to inversely correlate with NAFLD and other metabolic diseases, **Pfeuffer & Jaudszus (2016); Mika et al (2016); Yoo et al (2017)**. As demonstrated in our previous study that high dietary fat (HFD) intake reduces serum and liver OCFA, OCFA- producing gut microbiota and is associated with impaired liver lipid metabolism **Ampong et al (2021)**. Methionine adenosylated product, *S*-adenosylmethionine is a known substrate for methylation in epigenetic and epigenomic pathways, **Robinson et al (2016)**, and methylation has been linked to NAFLD directly and indirectly via PPARg. Likewise, *SCD1* also indirectly affected by promoter methylation of *LXR* in relation to NAFLD models Kim et al (2020). Hence, our current study sought to investigate the impact of LPD on OCFA and how methyl donor in LPD influences OCFA and other FAs. Accordingly, we observed a significant decrease in serum OCSFA and proportional decrease but not significant trend in the liver. In an attempt to see if the downregulation will be reversed after supplementing LPD with methyl donors, we observed upregulation of C17:0 and higher levels in other OCSFA where the changes were statistically insignificant in serum and liver. Methyl donors such as methionine, folate, betaine, and choline play important cellular functional roles including DNA methylation, phosphatidylcholine synthesis, and protein synthesis **Obeid (2013)**. The previous study reported that dietary methyl donors supplementation reduced fatty liver and modified the fatty acid synthase DNA methylation profile in rats fed an obesogenic diet **Cordero et al (2013)**.

In addition to the OCSFA dysregulation in protein restriction feeding, we also observed several saturated and unsaturated fatty acids including C10:0, C12:0, C14:0, C16:1, C16:0 C20:4n6 were altered in serum and liver in LPD fed mice and again these were all reversed in opposite direction when LPD was supplemented with methyl donors. This further suggests that methyl donors may have the potential to reverse the deleterious effect of “bad” fats whilst increasing the levels of “good” fats. The role of SFA, MUFA and PUFA in influencing NAFLD pathogenesis is well established **Juárez-Hernández et al (2015); Perdomo et al (2015); Properzi et al (2018)**.

A low protein diet feeding significantly upregulated the hepatic expression of *CD36* in mice relative to NPD. A previous study implicated *CD36* to play a crucial role in hepatic fatty acid uptake and hepatic steatosis in rodents, **Sheedfar et al (2014)**. *CD36* is a member of the class B scavenger receptor family with the ability to bind long-chain fatty acids, phospholipids, and collagen, **Febbraio et al (2002)**, and it’s less expressed in normal hepatocytes but its expression is increased with lipid-rich diets, hepatic steatosis, and NAFLD, **Wilson et al (2015)**. Whilst *FADS1* expression in the liver of mice fed LPD was not altered possibly reflecting the observation of PUFA profiles in the liver, we observed upregulation of this gene in MD-LPD fed mice suggesting that methyl donor lowers the risk of NAFLD. Previous studies have linked reduced function of D5D and decreased hepatic *FADS1* expression to be associated with NAFLD **Athinarayanan et al (2021)**. Catalase, an important antioxidant enzymes, is known to play key roles in lipid regulation. Its deficiency in mice has been reported to increase the likelihood of fatty liver **Heit et al (2017)**. Here, we observed an increase gene expression of *CATALASE* and *TRX1* in the liver of mice fed with methyl donor supplemented LPD. This means methyl donors are likely to guard against hepatic inflammation and fat accumulation.

In conclusion, the present study confirmed that OCFA and many novel metabolic pathways are dysregulated by consumption of a low protein diet and that these metabolic disturbances are reversed by methyl donor supplementation. In addition, several other fatty acids were persistently dysregulated during the development of NAFLD which were all reversed by methyl donor supplements. Therefore, this study provides the molecular mechanism in understanding the metabolic reprogramming in protein restriction-related metabolic disorders. Further interventional study designs to increase OCFA such as addition of branch-chain amino acids, dietary fish oils, propionate-enrich probiotics are required to improve metabolic health. Moreover, translational studies in human population involving methyl donor supplementation are needed to counteract the increasing prevalent of NAFLD and MS in developing countries.

## Acknowledgements

IA acknowledges the Commonwealth PhD scholarship, GHCS-2016–146 Commonwealth Scholarships Commission, UK, 2016–2019.

